# Genotypic diversity between surgical and nasal *Staphylococcus aureus* isolates

**DOI:** 10.1101/2020.05.27.120436

**Authors:** Dongzhu Ma, Patrick L. Maher, Kimberly M. Brothers, Nathan J. Phillips, Deborah Simonetti, A. William Pasculle, Anthony R. Richardson, Vaughn S. Cooper, Kenneth L. Urish

## Abstract

*Staphylococcus aureus* is a common organism in periprosthetic joint infection (PJI). Little is known about *S. aureus* genetic diversity in PJI as compared to nasal carriage. We hypothesized PJI *S. aureus* strains would be associated with increased virulence as compared to those from nasal carriage. Whole genome sequencing and multilocus sequence typing (MLST) was performed to genotype these two populations at high resolution. MLST revealed a variety of genotypes in both populations but many belonged to the most common clonal complexes. In nasal cultures, 69% of strains were of clonal complexes CC5, CC8, and CC30. In PJI cultures, only 51% could be classified in these common clonal complexes. Remaining strains were atypical, and these atypical strains in PJI were associated with poor host status and compromised immune conditions. Mutations in genes involved in fibronectin binding (*ebh, fnbA, clfA, clfB*) systematically distinguished later PJI isolates from the first PJI isolate from each patient. *S. aureus* isolated from nasal carriage and PJI specimens differ significantly, with the latter being more diverse. Strains associated with lower pathogenicity tended to be found in immunocompromised patients, suggesting the host immune system plays an important role in preventing PJI. Repeated mutations in *S. aureus* genes associated with extracellular matrix binding were identified suggesting an adaptive, parallel evolution in *S. aureus* during the development of PJI.

## Introduction

The primary organism associated with surgical site and implant infections is *Staphylococcus aureus*. It is a common commensal organism colonizing the nares and skin. As an opportunistic pathogen, it can be responsible for self-limited infections such as abscesses to life-threatening conditions such as sepsis, endocarditis, and pneumonia (1–4). These episodes of infection are self-limited and typically acute with either minimal or high morbidity and mortality (3). An exception includes surgical infection, especially those associated with medical devices. Surgical site infection is one of the most common health-care associated infections (5). Unlike most other *S. aureus* infections, surgical implant infections can be chronic, indolent, and challenging to treat.

Knee and hip arthroplasty or total joint replacement illustrate this problem. Arthroplasty, one of the largest major surgical procedures in the world by volume, has been named one the greatest medical innovation of the twentieth century (6). Total hip arthroplasty has a survival of approximately 60% at 25 years, an engineering marvel (7). The largest overall reason for failure of knee and hip implants is infection, termed periprosthetic joint infection (8–11). *S. aureus* is the most common organism associated with periprosthetic joint infection (12, 13).

*S. aureus* has a wide diversity between typical and atypical strains as demonstrated by the MLST types (14, 15). Little is known about this diversity relationship to pathogenesis. There is limited but strong evidence that PJI is related to host colonization (16). Similar to other types of surgical infection, much is known about host risk factors, but little is known about bacterial virulence factors associated with disease.

Given the lack of understanding behind the role of genetic diversity in *S. aureus* pathogenesis, we completed a prospective clinical study to compare the genetic diversity between nasal carriage and PJI clinical isolates. We hypothesized that clinical isolates from PJI would be associated with more virulent strains. Our hypothesis was incorrect. Compared to the expected clonal complex distribution in nasal carriage, *S. aureus* clinical isolates from periprosthetic joint infection had an increased association with atypical strains. There was an association between atypical strains and an immunosuppressed state of the host. There was a subset of patients with serial cultures during the infection process. In this group, we observed mutations in the surface attachment proteins that was associated with a phenotypic change in binding.

## Materials and Methods

### Study Design

A prospective observational study of patients diagnosed with PJI was performed. Data were acquired from electronic medical records, and institutional review board approval was obtained. The study was completed at a regional health system.

### Study Participants

The cohort was obtained by sequentially collecting clinical isolates from PJI cultures. Medical specimens for culture are sent to a central microbiology laboratory at our institution for microbiology culture. A clinical laboratory standard index (CLSI) protocol was used for specimen culture. *S. aureus* cultures from known PJI surgical cases were collected sequentially. Nasal carriage was collected by sequentially collecting nares cultures from the same hospital system. Inclusion criteria included *S. aureus* culture from a confirmed case of PJI. Musculoskeletal Infection Society (MSIS) Criteria (17) were followed with modified minor criteria including synovial nucleated white blood cell count threshold of 2,500 cells/μl and a synovial polymorphonuclear percentage greater than 65% as a secondary analysis of infection status (18–20). Any patient that did not meet this modified MSIS criteria was further excluded. *Staphylococcus aureus* samples were collected from our institution and grown in trypticase soy broth (TSB) contain 0.25% glucose, and then stored in 15% glycerol TSB medium at −80°C. A total of 98 isolates (47 PJI and 51 nares) were collected.

### Genomic bacterial DNA isolation

The MasterPure™ Gram Positive DNA Purification Kit (Lucigen Corp., USA) was used to isolate genomic DNA from *S. aureus* samples. Briefly, clinical *S. aureus* stock samples stored at −80°C were streaked out onto trypticase soy agar (TSA) plates and incubated overnight at 37°C. Single colonies were picked from TSA plates and inoculated into in 5 ml TSB medium and grown overnight at 37°C with 250 rpm shaking. One and one-half milliliter of overnight culture was pelleted by centrifugation and 150 μl of TE Buffer was added to resuspend the cell pellet and vortexed. One microliter of Ready-Lyse Lysozyme was added to each resuspended pellet of bacteria and incubated at 37°C until the bacterial cell wall was destroyed. Next, 150 μl of Proteinase K/Gram Positive Lysis Solution was added to the sample, mixed thoroughly and incubated at 65°C for 15 minutes, vortexing briefly every 5 minutes, then cooled to 37°C. The samples were chilled on ice for 5 minutes and 175 μl of MPC protein precipitation solution (Lucigen, USA) was added to 300 μl of the lysed sample and vortexed vigorously for 10 seconds. Cellular debris was pelleted by centrifugation at 4°C for 10 minutes at 12,000 × g. The pellet was discarded and the supernatant was removed and transferred to a clean microcentrifuge tube. 1 μl of RNase A (5 μg/μl) was added to each sample, mixed thoroughly and incubated at 37°C for 10 minutes. 500 μl of isopropanol was added to the recovered supernatant and tubes were inverted 30-40 times. Precipitated DNA was pelleted by centrifugation at 4°C for 10 minutes at 12,000 × g. Isopropanol was removed using a pipet tip without dislodging the DNA pellet. The pellet was rinsed with 70% ethanol and resuspended in 50 μl of TE Buffer.

### Fibronectin and Fibrinogen binding assay

*S. aureus* binding to fibronectin and fibrinogen was determined using a semi-quantitative adherence assay on 96-well tissue culture plates (Sigma-Aldrich, USA). Plates were coated with 100 μl of 0.02% sodium carbonate (pH 9.6) containing 2 μg/ml fibronectin (Sigma-Aldrich, USA) or 100 μl of phosphate-buffered saline (PBS) containing 5 μg/ml fibrinogen (Sigma-Aldrich, USA) and incubated overnight at 4°C. The plates were washed three times with PBS and then blocked with 100 μl of a 2% bovine serum albumin (BSA) solution for 1 h at 37°C. The wells were washed three times with 100 μl of PBS and 100 μl of bacteria (approximately 1×10^7^ cells) was added to the appropriate wells and incubated for 1 hour or 24 hours at 37°C. Wells were washed four times with 100 μl of PBS. Bacteria were fixed with 100 μl of 10% formaldehyde (Sigma-Aldrich, USA) for 10 minutes. 100 μl of 0.2% crystal violet (Sigma-Aldrich, USA) was added to each well for 10 minutes, cells were washed four times with distilled water and wells were air dried for 2 hours. 100 μl of 30% acetic acid (Fisher Scientific, USA) was added to dissolve the crystal violet and absorbance was measured at 570 nm using a Multiskan plate reader.

### Markerless ebh gene deletion using pKFT Vector

The markerless *ebh* gene deletion was developed from the protocol of Dr. Fuminori Kato (21). Briefly, a 1097-bp DNA fragment containing an upstream region of *ebh* was amplified from our clinical *Staphylococcus aureus* sample 1 genome DNA using primers *ebh* 5’-*Sal I* (5’- CCGGTCGACGTTGGCGTTGGATTTTCATCC-3’) and *ebh* 5’-*Xba I* (5’-TGATCTAGACTCGAGTGCTCCAGAATATAATAACAC-3’). The PCR product was digested with *Sal I* and *Xba* I, and subcloned into the same site of pKFT to obtain pKFT-*ebh*-up. Likewise, a 1067-bp DNA fragment containing a downstream region of *ebh* was amplified with primers *ebh* 3’-*Xba I* (5’-GCACTCGAGTCTAGATCAAAAACCTGCTGAATCACC-3’) and *ebh* 3’-*BamHI* (5’-CTCGGTACCGATGGATCCAACGGTATTCCCGAA-3’). The PCR product was digested with *Xba I* and *BamHI*, and subcloned into the same site of pKFT-*ebh*-up, yielding the allelic replacement vector pFK-*ebh*. The pFK-*ebh* plasmid was first transformed into DNA restriction system-deficient *S. aureus* RN4220, then the modified plasmid was isolated and electroporated into our clinical *S. aureus* sample 1 (This strain was tolerant to most antibiotics). Transformants were selected at 30°C on TSA plates containing tetracycline. Then, single colony transformants were grown at 30°C in 4 ml TSB containing tetracycline at 250 rpm shaking. Integration of the plasmid into the chromosome by a single crossover event was achieved by incubation at 42°C overnight on TSA plates containing tetracycline. Correct homologous recombination of the target region was verified by PCR using the primer set pUC-UV (5’- CGACGTTGTAAAACGACGGCCAGT-3’, plasmid) and *ebh* 5’-up (5’-GCAGGTGGTC TTGCTGATAA-3’, chromosome) or pUC-RV (5’-CATGGTCATAGCTGTTTCCTGTG-3’, plasmid) and *ebh* 3’-dn (5’-CTTCGTCGCCTCGATAGTATTT-3’, chromosome). Then, the correct integrants were grown at 25°C overnight with shaking in 10 ml TSB without any antibiotics; repeated twice. The cells were serially diluted and plated on TSA plates and incubated at 42°C. The excision of the plasmid region in the chromosome by a double-crossover event was screened for tetracycline-sensitive colonies by replica-plating candidates on TSA plates versus TSA plates containing tetracycline (3 μg/ml). Integrants were cultured overnight at 37°C. The markerless deletion mutants were screened by PCR using primers *ebh* 5’-inner (5’- TGCATCCGAGTGTTTGAAGTA-3’) and *ebh* 3’-inner (5’-CAACGACAGCTGAGCAATTAAG-3’) from tetracycline-sensitive colonies.

### Sequencing, Genome, and Bioinformatic Analyses

Genomic DNA was isolated from clones following overnight culture using a MasterPure™ Gram Positive DNA Purification Kit (Lucigen Corp., USA). The sequencing library was prepared using the Illumina Nextera kit following published methods (22, 23). All samples were sequenced to at least 150-fold average coverage using an Illumina NextSeq 500. Following sequencing, we trimmed the samples with Trimmomatic (version 0.36) (24) and assembled the genome of the first isolate using SPADeS (25) and then annotated this initial genome using Prokka (26). All genomes were subsequently compared by mapping reads from later isolates against this initial assembled reference using breseq (version 0.28) (27). Mutation calls were manually curated to remove false positives due to reference assembly error, misaligned reads, and to consolidate sequential single base pair insertions or deletions into a single multiple base pair insertion or deletion.

### Data access

All sequenced genomes are available as BioProject PRJNA628522 in NCBI.

### Statistical Analysis

Demographic characteristics of our sample participants were analyzed using univariate descriptive statistics. Means and standard deviations were calculated for approximately normally distributed variables; medians and inter-quartile ranges (IQRs) were computed for continuous variables with skewed distributions. Frequencies and percentages were determined for categorical variables. Similar univariate techniques were used to characterize secondary outcomes such as follow-up, mortality and secondary surgical procedures.

## Results

### PJI infections are more frequently caused by atypical S. aureus lineages

Multilocus sequence typing (MLST) is a molecular technique that characterizes microbial species using the sequences of seven house-keeping genes. The alleles of the seven loci define the sequence type (ST) of each isolate (28). Using MLST to initially compare *S. aureus* diversity between PJI isolates and nasal carriage, the PJI isolates traced to more diverse lineages than those from nasal carriage, despite the majority belonging to common clonal complexes. In nasal cultures, 69% of strains were of clonal complexes CC5, CC8, and CC30, and all MRSA strains were of CC5 and CC8 (Figure 1A). In PJI cultures, only 51% could be classified in these common clonal complexes (Figure 1B). There was a strong trend in a larger proportion of strains from atypical clonal complexes in the PJI cohort as compared to the nasal carriage cohort (p = 0.076) (Figure 1C).

**Figure 1.**
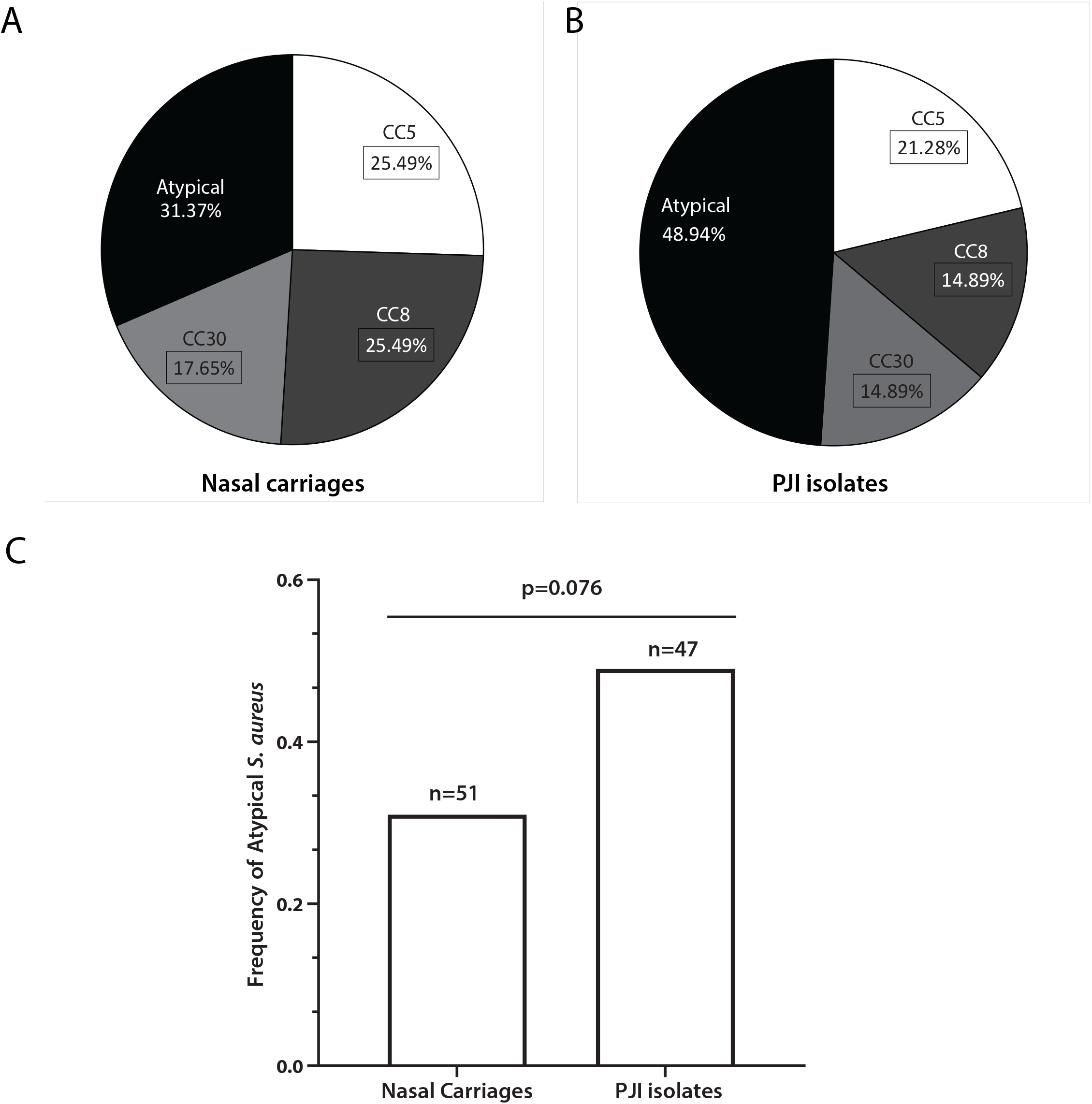
*S. aureus* diversity between PJI isolates and nasal carriages. A total of 98 isolates were analyzed (47 PJI and 51 nares). MLST classification was used to analyze *S. aureus* diversity between PJI isolates and nasal carriage. (A) Clonal complexes of *S. aureus* lineages from nasal carriages. (B) Clonal complexes of *S. aureus* lineages from PJI isolates. (C) PJI strains have a larger proportion of atypical *S. aureus* lineages as compared to isolates from the nares.

Atypical strains are usually less pathogenic than more commonly observed, “epidemic” clonal complexes in *S. aureus* infection (29, 30). If atypical strains were less pathogenic, we reasoned these atypical infections should be associated with poor host status and compromised immune conditions. Many disease processes are associated with immune compromised status including diabetes mellitus, smoking, autoimmune disease, and immune suppressive therapy including steroid use (31–33). When the proportion of atypical *S. aureus* strains in PJI patients associated with these conditions were compared to PJI patients without these conditions, these atypical strains were associated with poor host status and compromised immune conditions (p < 0.01, Figure 2).

**Figure 2.**
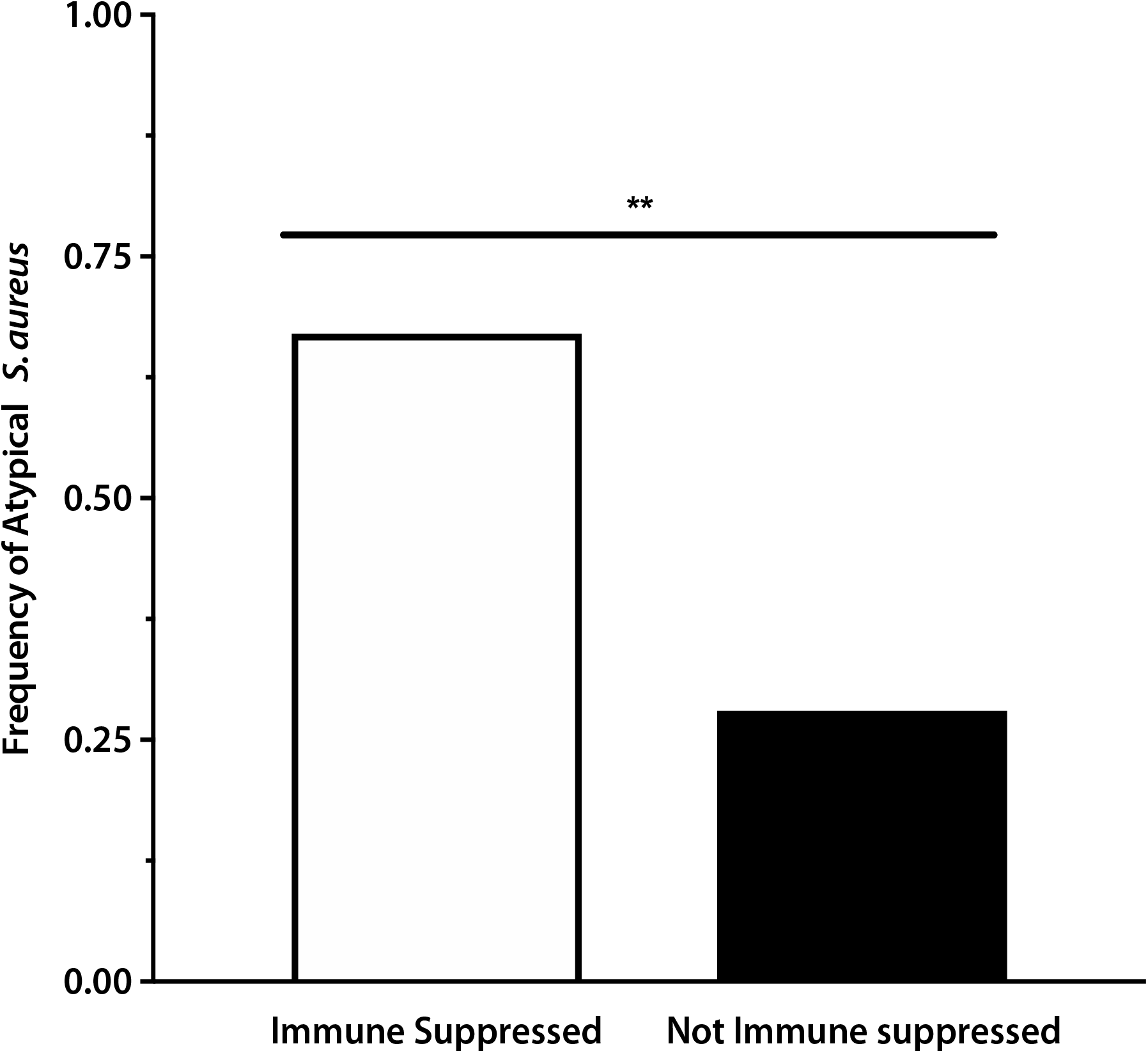
Frequency of atypical *S. aureus* in immune compromised patients Immune status is associated with many disease processes. When compared to the proportion of atypical *S. aureus* strains in immune compromised patients to those without, the immune suppressed patients have higher frequency of atypical *S. aureus* lineages. ** p<0.01.

### Mutations in fibronectin binding genes occurs in subsequent/latter PJI isolates

Observing an association between atypical strains and PJI was unexpected. If atypical strains are not associated with common *S. aureus* infections, then this suggested that the local environment in PJI had unique characteristics that allowed *S. aureus* to thrive. We hypothesized that selection on these infecting populations would enrich mutations in traits adaptive in this foreign environment. To test this, we quantified genetic diversity at a local level. In the PJI cohort, there were a series of 4 patients with isolates collected at multiple locations and time points. *S. aureus* infections are known to originate from a small clonal population (34). If there was little genetic diversity, then we expected these populations from each patient to remain clonal. Whole genome sequencing demonstrated that multiple isolates from the same patient were nearly identical allowing mutations to be identified. An array of mutations were identified in the series of isolates from each patient (Figure 3). Mutations involved in fibronectin binding (*ebh, fnbA, clfA, clfB*) systematically distinguished later PJI isolates from the first PJI isolate from each patient. The mutations primarily occurred outside the normal reading frame nonsynonymous single nucleotide polymorphisms (SNPs) (Supplemental Figure 1). One of the isolates upon initial infection in synovial fluid had mutations in *ebh*, lysostaphin, multidrug resistance, pheromone binding and epimerase. After initial treatment, there was a recurrence of the infection in the same knee 90 days later. Upon reinfection the new isolate only had mutations in *ebh* and lysostaphin (Figure 3C). This implied the strain was present in the initial infection, not isolated during the initial infection, and gave rise to sample 2 and 33. As this isolate was not isolated during the initial infection, we could not determine what conditions gave rise to a loss of the other genetic mutations found in initial infection. The later isolates cultured from synovial fluid had the same genotype as this initial uncultured isolate (Figure 3C).

**Figure 3.**
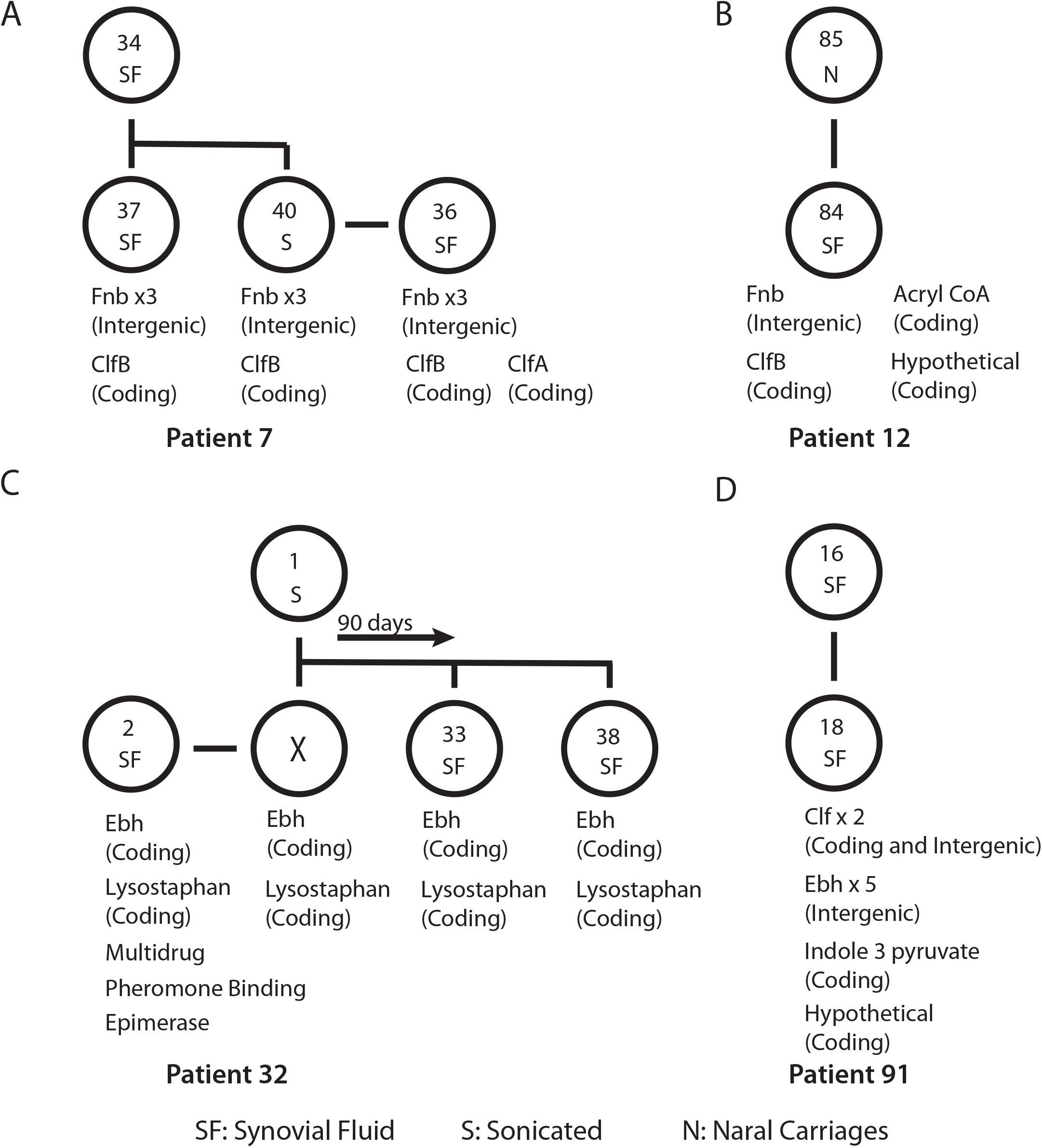
Mutations identified in different *S. aureus* isolates from the same patient were primarily involved with bacteria binding There were 4 patients that had multiple isolates collected through the infection process. Whole genome sequencing identified mutations involved in fibronectin binding (*ebh, fnbA, clfA, clfB*) that systematically distinguished later PJI isolates from the first PJI isolate from each patient. (A) Mutations from a series of clinical isolates in patient 7. (B) Mutations were identified between the nares and PJI isolate of patient 12. (C) Mutations from a series of clinical isolates in patient 32. In this case, after treatment there was a recurrence of infection at 90 days. Mutations were also found in lysostaphin, multidrug resistance, pheromone binding and epimerase. (D) Mutations from a series of clinical isolates in patient 91. SF: Synovial Fluid; S: Sonicated; N: Nasal Carriages.

### *MSCRAMM binding is altered in ebh* mutants

We questioned if these mutations conferred an observed phenotype. As the majority of the mutations involve MSCRAMM binding, we quantified the fibronectin and fibrinogen binding of each isolate and compared it to the initial strain (Figure 4A and B). Each isolate had a variation in fibronectin and fibrinogen binding as compared to the parent isolate. One patient had isolates from multiple extended time points from a recurrent infection (Figure 3C). Mutations in *ebh* were common. To test if this mutation resulted in an observed phenotype, we created an *ebh* mutant strain from the initial clinical isolate. The *ebh* mutant had significantly (p < 0.01) lower fibronectin and fibrinogen binding (Figure 4A and B) as compared to the clinical isolates. We confirmed the mutation with PCR (Supplemental Figure 2). Ebh is the largest gene in the *S. aureus* genome at 1,1000 KD (35), preventing our ability to complete complement analysis. As an alternative method, we completed a whole genome sequence of the *ebh* clinical isolate mutant to detect any other differences. No other mutations were identified.

**Figure 4.**
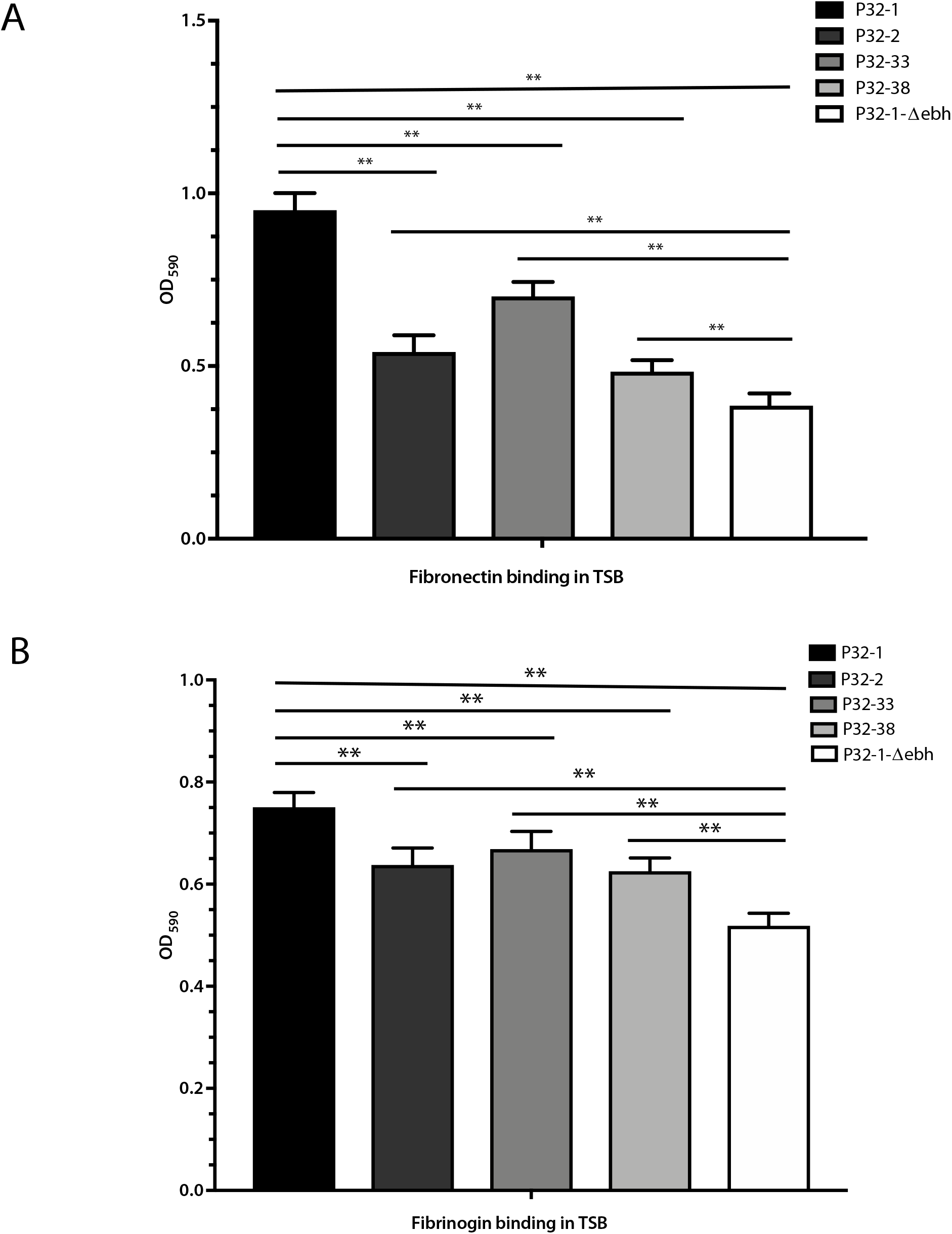
Phenotype changes in fibronectin and fibrinogen binding between clinical isolates with genotype changes in *ebh*. The fibronectin and fibrinogen binding of each isolate from one patient were compared to their initial strain as measured by crystal violet (CV) absorption assay in 96 well plates coated with fibronectin or fibrinogen. The results demonstrated that fibronectin (A) and fibrinogen binding (B) was altered in *ebh* mutants. The *ebh* mutant had lower fibronectin and fibrinogen binding (A and B) as compared to all of the clinical isolates. ** p<0.01

## Discussion

The role of bacteria genetic diversity in the pathogenesis of surgical infection is poorly understood. We selected total knee arthroplasty periprosthetic joint infection (PJI) as a model since it serves as a valuable example and clinical model for this problem. The objective of this study was to compare global and local diversity in *S. aureus* associated with PJI. *S. aureus* isolated from PJI had an association with atypical MLST strains as compared to nasal carriage. This was unexpected as common MLST strains are thought to be more pathogenic. These atypical strains were more diverse and less virulent in comparison to nasal carriers. The presence of a foreign body (i.e. prosthesis) in the joint could be a possible explanation for these differences. In addition, our findings from this study demonstrate an increased prevalence of atypical strains in immunosuppressed patients providing further evidence that PJIs are associated with poor host status. At a local level, we observed evidence of adaptive mutations and selective pressure in these different environments.

*S. aureus* is a human commensal frequently associated with human skin and mucosal surfaces. In health-care associated MRSA infections CC5, CC8, and CC30 are most common lineages found in North America, while in South America it is predominantly CC5 and CC8 (36, 37). This is in contrast to Europe and Asia where the distribution is CC5, CC8, and CC22 lineages (36, 37). The primary reservoir of *S. aureus* is the anterior nares, and the rates of *S. aureus* nasal carriage can be up to 59% (2, 38, 39). Nasal carriage of *S. aureus* is a risk factor in various surgical infections (2, 38, 39). Bacteremia and surgical infection are often thought to originate from this anatomic location (16, 40–42). There are about 27-84% of *S. aureus* surgical site infections identified with isolated strains originating from the nose (43). At an epidemiologic level, we observed a large difference in genetic diversity between nasal carriage and surgical infection in PJI. Our results support previous observations that the most common isolates identified in the nose consist of primarily the CC5, CC8, and CC30 lineages (44–47). Melles reported that CC5 and CC8 account for 43.8% of the isolates from healthy humans, and CC30 accounts for 47.3% (48). This is the first larger observational report of *S. aureus* genetic diversity observed in PJI. We observed a greater than expected population of atypical strains in PJI that was more diverse than in nasal carriage. We cannot comment on direct comparisons in a single host between nasal carriage and surgical infection. Our study was not designed to look for self-transmission. In order to capture transitory nasal carriage a prospective longitudinal study would be needed to assess the origins of *S. aureus* in PJI. A study comparing the *S. aureus* isolates between nasal carriage and surgical infection site showed a similar pattern of *S. aureus* clonal complexes in six patients, which indicated that the surgical site infection in these cases likely originated from the host (16).

Gram-positive bacteria express a variety of virulence factors, which include surface proteins that are crucial for bacterial colonization and infection. Microbial surface components recognizing adhesive matrix molecules (MSCRAMMs) are a type of cell wall anchored protein that consist of IgG-like folded arrays that participate in ligand binding (49). In *S. aureus*, the MSCRAMMs that are present are used to attach to proteins on the host tissue, including human fibrin and fibronectin. Otto et al. demonstrated that fibrin and fibrinogen concentrations are much higher in the synovial fluid allowing *S. aureus* to persist as large aggregate biofilms through MSCRAMM binding (49, 50). In our study, increased genotype variations in MSCRAMMs (*ebh, fnbA, clfA,* and *clfB*) were observed in the later PJI isolates. These genetic mutations were associated with an observed change in binding phenotype of fibronectin and fibrinogen indicating an overall selection pressure to alter *S. aureus* binding and adhesion. Mutation of surface attachment protein genes from independent ancestors suggests an adaptive parallel evolution in *S. aureus* during development of PJI.

Our PJI studies identified genotype variation in MSCRAMM binding proteins in later patient isolates. Since the most common mutations were found in binding proteins, our group focused on *ebh*, a highly conserved 1,100-kDa protein, which has homology to other ECM binding proteins (35). *Ebh* resides on the bacterial surface and forms a specialized structure to stabilize the cell wall against osmotic stresses and is involved in cellular adhesion (35). Mutations in *ebh* result in a weakened peptidoglycan, an irregularly shaped cell wall, and a much larger bacterial cell volume (51). *Ebh* facilitates *S. aureus* binding to fibronectin through FnbA and FnbB, and plays an important role in biofilm formation working together with polysaccharide intracellular adhesion (PIA) and extracellular DNA (52–55). Mutations of *ebh* in *S. aureus* are unable to withstand changes in osmotic pressure (52) and show diminished virulence in mice (51). Clinical isolates of *S. aureus* exhibit different fibronectin adherence (56). In our study, mutations involving fibronectin binding distinguished later PJI isolates from the initial PJI isolate from each patient. We focused on *ebh* since it is the largest MSCRAMM binding protein found in *S. aureus*. Mutational analysis of *ebh* revealed a significant reduction in fibronectin binding suggesting the importance of *ebh* in establishing a biofilm in PJI. Our results suggest there may be a subtle adaptive selection for *ebh* binding.

There were some limitations to this study. First, it was unknown if the infection was from a single clone or polyclonal. Only one clone was picked for genome analysis from each patient. For most cases, infections are believed to originate from a single clone (57). Genotype changes observed between different isolates from the same patient were not missense or nonsense mutations, but primarily with SNPs. Importantly, we confirmed at the level of the organism that at least one of these mutations were associated with a phenotype.

There is a large difference in diversity between *S. aureus* isolated from nasal carriage and PJI specimens. *S. aureus* associated with PJI was more diverse with a higher proportion of atypical strains. These atypical strains are associated with lower pathogenicity. The frequency of these atypical strains were higher in immune suppressed patients versus healthy suggesting the host immune system plays an important role in preventing PJI. Repeated mutations in *S. aureus* genes associated with extracellular matrix binding were identified suggesting an adaptive, parallel evolution in *S. aureus* during the development of PJI.

---

Figure S1. *S. aureus* isolates targeted area sequence alignment. Single nucleotide polymorphisms (SNPs) of target gene areas from different patient isolates.

Figure S2. *ebh* deletion from clinical isolate

Schematic diagram for *ebh* deletion mutation from clinical isolate 32. (A) pKFT-*ebh* shuttle vector used in the *ebh* deletion mutation procedure. (B) Detail of the *ebh* fragments used in the pKFT-*ebh* vector were demonstrated in USA300-FPR3757 (JE2) strain background. (C) Confirmation of *ebh* deletion by PCR method. * indicates correct colonies with *ebh* deletion.

